# Circadian Rhythm Analysis Using Wearable Device Data: A Novel Penalized Machine Learning Approach

**DOI:** 10.1101/2020.02.20.957076

**Authors:** Xinyue Li, Michael Kane, Yunting Zhang, Wanqi Sun, Yuanjin Song, Shumei Dong, Qingmin Lin, Qi Zhu, Fan Jiang, Hongyu Zhao

**Author notes:** **Address correspondence to:** Hongyu Zhao, PhD, Department of Biostatistics, Yale School of Public Health, New Haven, CT, USA, Postal address: 60 College Street, Suite 201, New Haven, CT, USA.

## Abstract

**Background:** Wearable devices have been widely used in clinical studies to study daily activity patterns, but the analysis remains the major obstacle for researchers.

**Study Objective:** This study proposed a novel method to characterize sleep-activity rhythms using actigraphy and further used it to describe early childhood daily rhythm formation and examine its association with physical development.

**Methods:** We developed a machine learning-based Penalized Multi-band Learning (PML) algorithm to sequentially infer dominant periodicities based on Fast Fourier Transform (FFT) and further characterize daily rhythms. We implemented and applied the algorithm to Actiwatch data collected from a 262 healthy infant cohort at 6-, 12-, 18-, and 24-month old, with 159, 101, 111, and 141 subjects participating at each time point respectively. Autocorrelation analysis and Fisher’s test for harmonic analysis with Bonferroni correction were applied to compare with PML. The association between activity rhythm features and early childhood motor development, assessed by Peabody Developmental Motor Scales-Second Edition (PDMS-2), was studied through linear regression.

**Results:** PML results showed that 1-day periodicity is most dominant at 6 and 12 months, whereas 1-day, 1/3-day, and 1/2-day periodicities are most dominant at 18 and 24 months. These periodicities are all significant in Fisher’s test, with 1/4-day periodicity also significant at 12 months. Autocorrelation effectively detected 1-day periodicity but not others. At 6 months, PDMS-2 is associated with assessment seasons. At 12 months, PDMS-2 is associated with seasons and FFT signals at 1/3-day periodicity (*P*<.001) and 1/2-day periodicity (*P*=.04). In particular, subcategories of stationary, locomotion, and gross motor are associated with FFT signals at 1/3-day periodicity (*P*<.001).

**Conclusions:** The proposed PML algorithm can effectively conduct circadian rhythm analysis using time-series wearable device data. Application of the method effectively characterized sleep-wake rhythm development and identified the association between daily rhythm formation and motor development during early childhood.

## Introduction

Wearable devices have been increasingly used in research recently, as they can provide continuous objective monitoring of activities as well as vital sign data such as body temperature and pulse rates[1-3]. In sleep and activity studies, researchers are focused on the generated actigraphy from wearable device activity data to study sleep and activity patterns as an alternative to sleep diaries and polysomnography (PSG) [1, 4]. The device usually uses an accelerometer that works by monitoring acceleration in one or more directions, and this wristwatch-like device is often worn on the wrist to record activity continuously for several days. Either the raw data or the transformed activity count data can be used to study sleep-wake patterns and screen sleep disorders [4, 5]. Actigraphy not only avoids the subjectivity and bias issues with sleep diaries but also overcomes the drawbacks of PSG, such as high costs, in-lab setting, intrusive measures, and difficulty in long-time monitoring.

Continuous objective monitoring by the wearable device provides researchers with the opportunity to conduct circadian rhythm studies. Circadian rhythms are endogenous and entrainable biological processes that follow a period of approximately 24 hours, and many physiological phenomena such as sleep-wake patterns, body temperature, and hormone levels all exhibit circadian rhythms. For humans, most circadian rhythms are under the control of the pacemaker located in the suprachiasmatic nuclei in the anterior hypothalamus of the central nervous system, and suprachiasmatic nuclei accepts environmental information such as the light/dark cycle to adjust the 24-hour cycle [6]. However, 24-hour human circadian rhythms are not mature at birth, when the predominant rhythm is ultradian, and the circadian rhythms of sleep-wake cycles and body temperature gradually develop during the first year after birth [6-9]. Many studies have investigated how circadian rhythms develop through childhood into adolescence and adults, and how they are related to health issues such as sleep problems, mental problems, and disease risks, just to name a few [8, 10-14]. It is noteworthy that the development of circadian rhythms during early childhood is associated with disease risk factors and can affect both childhood and adult life [8]. Therefore, it is important to conduct circadian rhythm studies to get a thorough understanding of the formation and consolidation of daily activity rhythms during early development as well as the association between changes in daily rhythms and health conditions.

Actigraphy generated from wearable devices has been validated to provide reliable information on sleep and circadian rhythms [15]. However, the analysis of time series data from actigraphy remains the major obstacle for researchers. Current major statistical methods are either parametric based on cosinor analysis or nonparametric [16-21]. These methods do not specifically focus on periodic information and are not specifically suitable for populations whose sleep-wake rhythms are not sinusoidal, such as circadian disorder patients, or not mature, such as young infants and toddlers [6-10, 12]. Therefore, traditional approaches targeting normal daily rhythms might not work, as detailed activity rhythms cannot be captured. There is a need to develop appropriate methodology to extract periodic information and study detailed circadian patterns of all populations to better characterize daily rhythms.

Here we propose a Penalized Multi-band Learning (PML) approach that can complement current methods to characterize daily rhythms based on periodic information in time series wearable device data. PML extracts periodic information using Fast Fourier Transform (FFT) and then performs penalized selection based on regularization, a classic approach used in machine learning, to identify dominant periodicities and further characterize daily rhythms [22, 23]. In this paper, we first present the proposed PML approach in details and discuss its usefulness and advantages compared to other methods. Then, we present one application of the method to early childhood wearable device activity data, in which we were able to characterize the formation and consolidation of sleep-activity rhythms and further studied its association with physical development during early childhood.

## Methods

### Data

The study subjects are from a 262 healthy newborn cohort recruited in 2012-2013 by Shanghai Children’s Medical Center, Shanghai, China. Actiwatch data were collected at 6, 12, 18, and 24 months old, with 159, 101, 111, and 141 subjects at each time point respectively, and not all subjects from the cohort participated each time. Infants and toddlers were required to wear Actiwatch-2 (Phillips Respironics Mini-Mitter) on the ankle for seven consecutive days. Wearing on the ankle is commonly recommended for young infants/toddlers [24]. The Actiwatch-2 utilizes a piezoelectric sensor to detect accelerations between 0.5 and 2.0g with a frequency response range between 0.35-7.5Hz, and activity counts summarize accelerations over each epoch. The data output format for Actiwatch-2 was configured to be activity count per 1-minute epoch. Based on sleep diaries and activity plots for each individual, days showing non-wear periods with straight lines of zero activity counts were removed. Non-wear periods can be differentiated from sedentary behaviors or sleep as the former gives almost all zeros while the latter gives non-zero activity counts every now and then. Figure 1 shows the activity plots for subject ID 17, and it can be seen that at 6 months, low and high activities are intermittent during the day, suggesting multiple daytime naps, while near-zero activity levels at night suggests long nighttime sleep. At 12 months, three activity peaks, one morning nap and one afternoon nap can be identified. Then at 18 and 24 months, two activity peaks formed and stabilized, showing one afternoon nap only. The daily activity rhythm developed and stabilized as the infant grew. In addition to Actiwatch data, demographic information and family information were collected at baseline, such as child sex, child birthdate, parental age, child birth weight and body length, parents’ heights and weights, parental educational levels and working statuses, and family income. Peabody Developmental Motor Scales-Second Edition (PDMS-2) were used to assess early childhood physical development at 6, 12, 18, and 24 months [25]. The institutional review board of the Shanghai Children’s Medical Center, Shanghai Jiao Tong University approved the study and the approval number is SCMCIRB-2012033. Parents of children who participated in the study all gave written informed consent.

**Figure 1.**
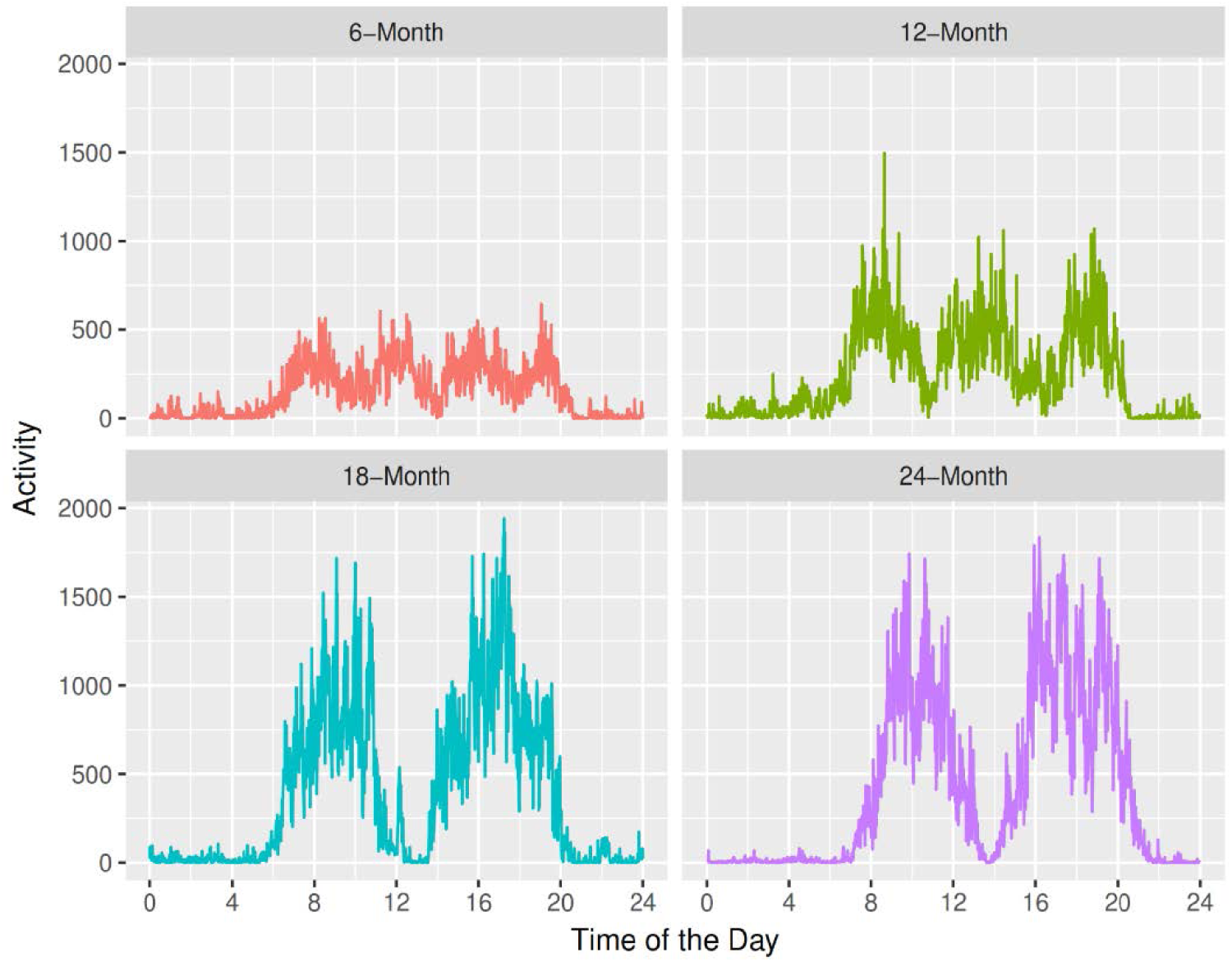
The activity plots for ID 17 at 6, 12, 18, and 24 months respectively, with activity counts averaged across seven days at each time point.

### Fast Fourier Transform

To describe the consolidation of sleep-activity rhythms during early childhood, we utilize periodic information to characterize daily rhythms. Specifically, we use Fast Fourier Transform (FFT) to convert time-domain signals into frequency-domain spectrum in order to extract periodic information. We analyze the original data to allow for non-24-hour sleep-wake rhythm detection. Figure 2 shows FFT results for ID 17 at each age respectively. 1-day periodicity is most dominant at all time points. 1/5-day and 1/4-day periodicities can be identified at 12 months. 1/2-day and 1/3-day periodicities did not become dominant until 18 and 24 months. It is noteworthy that each periodicity is not interpreted alone but combined to understand the overall pattern. As suggested on the right panel of Figure 2, the combined 1/5-day and 1/4-day periodicities form the three-peak two-nap pattern at 12 months. Likewise, the combined 1/2-day and 1/3-day periodicities exhibit the two-peak one-nap pattern at 18 and 24 months. Therefore, the combination of dominant periodicities can be utilized to capture main sleep-activity patterns at each age and describe the gradual consolidation of daily rhythms in early childhood development.

**Figure 2.**
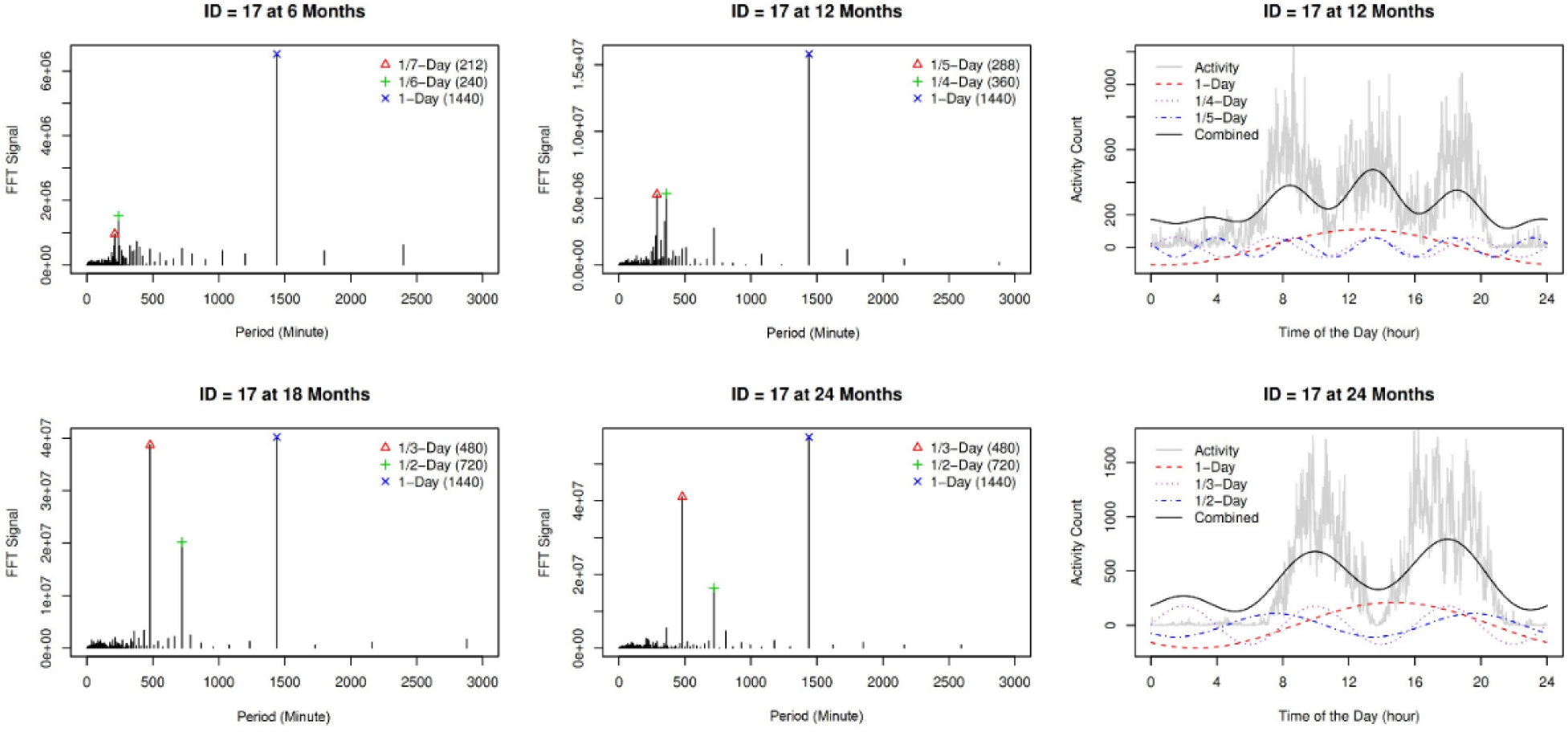
Left panel: Fast Fourier Transform (FFT) results for ID 17 at 6, 12, 18, and 24 months, respectively. Right panel: top three periodicities and the combined plotted on 1-day observation for ID 17 at 12 months and 24 months, respectively.

### Identification of Dominant Periodicities

The PML algorithm is as follows. Let matrix *X* ∈ *R*^*n*× *p*^ denote FFT results, where *n* denotes the number of individual observations, and *p* denotes the number of periodicities from FFT. Specifically, *X* = (*x*_1_, *x*_2_, …, *x*_*p*_), where *x*_*j*_ is the vector of length *n* for the *j*th periodicity.

Let Θ be the diagonal matrix selecting columns from *X* such that 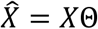 and 0 ≤*θ*_*j,j*_≤1, *j* =1,…, *p*:

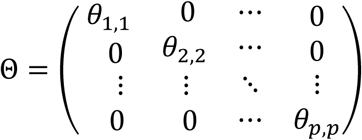

Θ identifies columns of dominant periodicities from *X* in the way that dominant periodicities corresponding to nonzero *θ*_*j,j*_′*s* are selected. We minimize the squared Frobenius norm 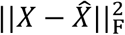, the sum of the squared elements of the matrix. Using properties of the Frobenius norm, we can get:

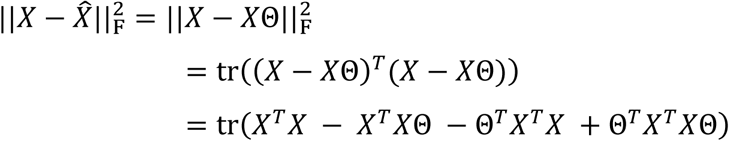

Because *X*^*T*^*X* is fixed, it is equivalent to minimize:

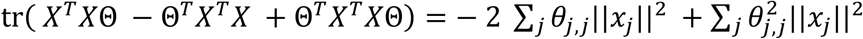

In order to estimate Θ and identify dominant periodicities, we use a penalized selection method similar to Lasso, a widely used method in shrinkage and selection of a subset of features in regression models and machine learning approaches [23]. In regression, Lasso penalty is most effective in selecting a few important features while suppressing regression coefficients of other non-selected features to 0 [23]. In our case, Lasso penalty serves to select a few dominant periodicities through diagonal elements of Θ instead of regression coefficients. Further, we add an elastic-net like penalty term onto the Frobenius norm, namely a combination of L1 and L2 norms[22]:

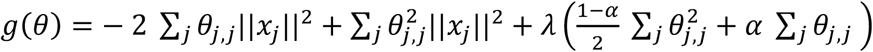

where *λ* is the tuning parameter and *α* controls the balance between the L1 and L2 norms. Note that *θ*_*j,j*_’s are nonnegative and thus we do not need to take the absolute value for the L1 norm. By setting *λ* large enough, all diagonal elements of Θ, namely all *θ*_*j,j*_′*s*, are suppressed to zero and no periodicities are selected. As *λ* decreases, some *θ*_*j,j*_′*s* become nonzero and they correspond to the most dominant periodicities that are selected sequentially according to how dominant they are.

To minimize *g*(*θ*), we take the partial derivative of *g*(*θ*) with respect to each 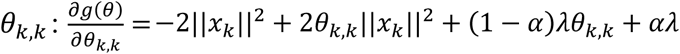, which is convex and also subject to the constraint 0 ≤ *θ*_*k,k*_ ≤ 1. Thus, we have:

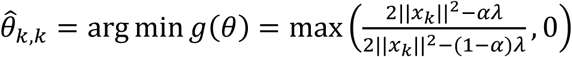

If we only have the L1 penalty, *α* = 1 and 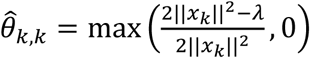. In our case, we use Lasso L1 penalty alone and train *λ*, because we want to select the most important periodicities while suppressing other periodicities to 0. However, we still keep the L2 norm in the original model as an option as it might be helpful in future tasks, such as prediction, classification, reconstruction of curves, and so on.

We use mean squared error (MSE), which is equivalent to the squared Frobenius norm 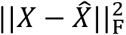, as the criterion for choosing *λ* and the number of nonzero *θ*_*j,j*_′*s* (the number of dominant periodicities selected) as well as evaluating how much variability is not explained by the selected periodicities. We did not choose cross-validation because results showed that the test dataset error curve is monotonous. We train *λ* from 2 · 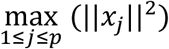 to 0, as 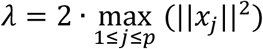 suppresses all *θ*_*j,j*_’s to 0 and *λ* = 0 gives no penalty. By decreasing *λ*, we identify dominant periodicities sequentially and characterize the daily sleep-activity rhythm at each age. An R package named PML has been developed (https://CRAN.R-project.org/package=PML) for the implementation of the PML algorithm [26].

### Comparison with Other Methods

To rigorously conduct statistical tests and select significant periodicities, we apply Fisher’s Test for Harmonic Analysis [27]. It is a sequential test for ordered statistics, and periodicities are first ordered and then tested for significance. If one periodicity is statistically significant, the next one will be tested further. If not, the sequential test will stop. At each step, the critical value at which to declare statistical significance is different. In some publications the method may not be implemented correctly so we included the sequential test in the R package for easy implementation. Because we are performing multiple testing, Bonferroni correction is used to adjust p-values. If we conduct the tests at significance level *α*, we will reject the null hypothesis if p-value ≤ *α*/*p*, where *p* is the number of periodicities, and conclude that the periodicity is significant.

To evaluate the effectiveness of the PML algorithm, we compare it with the standard approach autocorrelation. Autocorrelation *r*_*k*_ is calculated between activity measurements with a time lag *k*, and the coefficient *r*_24_ for a 24-hour time lag is of primary interest in circadian studies [28]. *r*_*k*_ ranges between −1 and 1, and a *k*-hour periodic pattern can show a higher value of correlation coefficient *r*_*k*_. In the plot of *r*_*k*_ against the time lag *k*, a peak around *k*=24 can be observed when there is a dominant circadian pattern of 24-hour periodicity. We plot autocorrelation against the time lag to compare the autocorrelation method with our algorithm.

### Association between Daily Rhythms and Motor Development

We further conduct linear regression to study the association between the consolidation of daily activity rhythms and early childhood physical development. PDMS-2 is considered as early childhood developmental assessment and used as the outcome. If PDMS-2 total motor standard scores are found to be associated with daily rhythm features, the standard scores for subtests including stationary, locomotor, object manipulation, grasping and visual motor integration as well as gross motor and fine motor are used as the outcome to further examine which specific subcategory is associated. Gross motor represents the overall performance on stationary, locomotion and reflexes (6 months) or object manipulation (12, 18, 24 months) for infants, and fine motor represents the overall performance on grasping and visual-motor integration.

The FFT signals at dominant periodicities identified by PML are used as daily rhythm features and considered as covariates in the model. In addition to periodic features, demographic information and family information as potential confounders are also considered in the model. Backward selection is used in the model fitting process. While some variable (denoted as variable A here) may appear to be statistically significant in the complete model, after removal of insignificant variables in the variable selection process, variable A may become insignificant. In such cases, variable A will also be removed in the final model to achieve parsimony.

Linear regression is conducted at 6, 12, 18, 24 months respectively to study the association between daily rhythms and motor development. For final model comparison, *r*^2^ that measures the proportion of the variation in the outcome explained by the model and also the adjusted *r*^2^ that modifies *r*^2^ based on the number of predictor variables were also calculated. All statistical analyses were conducted in R (Version 3.3.2).

## Results

### Identification of Dominant Periodicities

As shown in Figure 3, at each age, we plot MSE against the number of nonzero *θ*’s, and specifically, we only plot the points where the number of nonzero *θ*’s (periodicities selected) increases as the penalty *λ* decreases. For 6 months and 12 months, the first harmonic at 1-day is most dominant, as we can observe a large dip in MSE when the first periodicity is selected while the following periodicities that are further selected do not cause the same level of decrease. For 18 and 24 months, the first three periodicities of 1-day, 1/3-day, 1/2-day are most dominant, as selecting the first three can lead to relatively large decrease in MSE. The results indicate that during the first year, only 1-day periodicity is formed and stabilized in the infant population. Not until 18 months is the sleep-activity rhythm stabilized, showing the pattern of one nighttime sleep, one daytime nap, and two daytime activity peaks.

**Figure 3.**
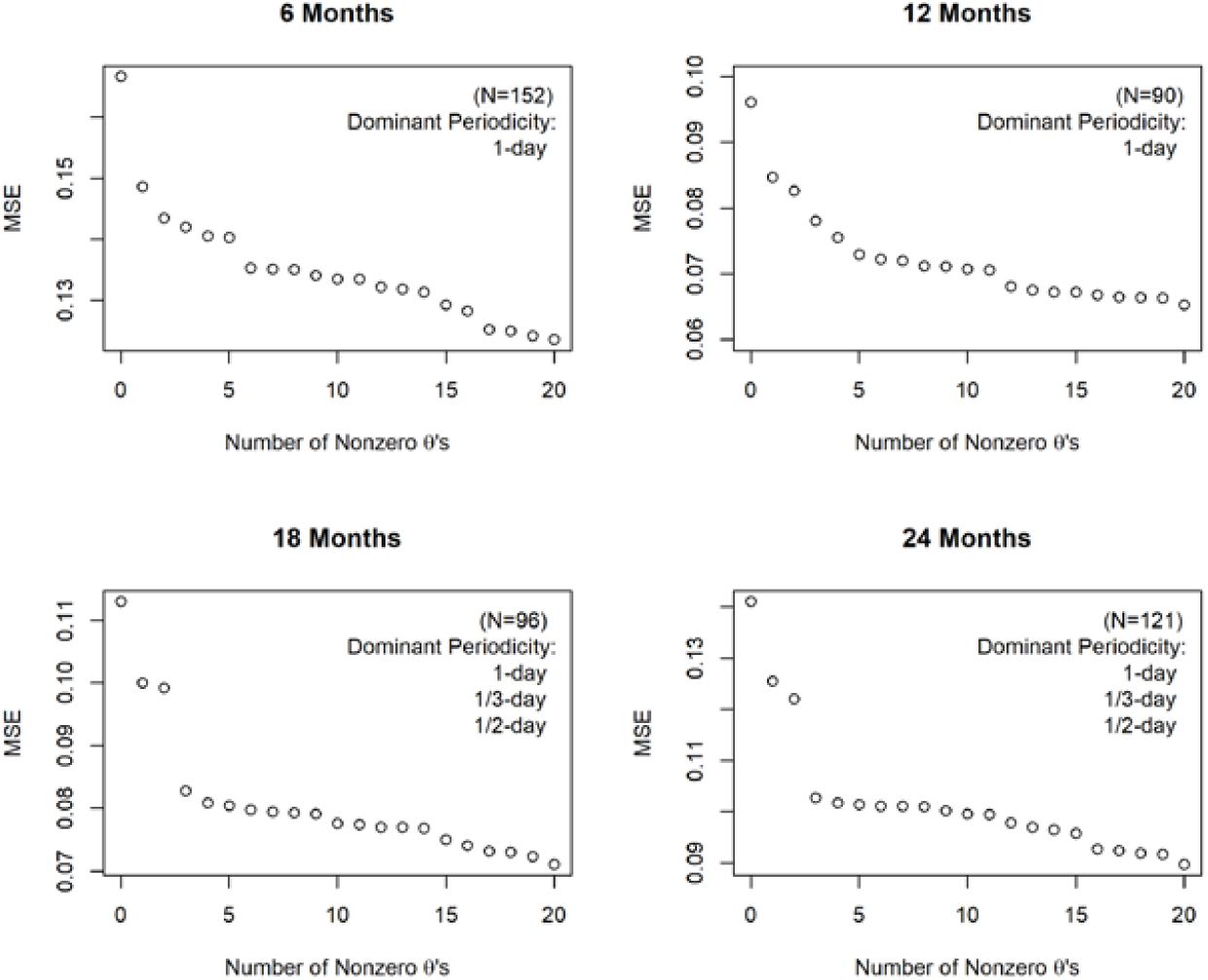
Mean squared error (MSE) against the number of nonzero *θ*’s as the penalty term *λ* decreases, at 6, 12, 18, and 24 months, respectively.

Fisher’s tests give similar results. As shown in Table 1, only 1-day periodicity is significant at 6 months (*P* < 10^−5^), because for infants at this stage, sleep-activity patterns have already adjusted to a 24-hour cycle. However, daytime activities have not been stabilized yet and variations exist across days. Then at 12 months, four periodicities are significant (*P* < 10^−5^). It is because infants’ sleep-activity patterns start to stabilize, but there are variations across individuals: some take one nap in the afternoon while others take two naps, one in the morning and one in the afternoon. The one-nap pattern can be captured by the 1/3-day periodicity, while the two-nap pattern can be captured by the 1/4-day periodicity, as previously shown in Figure 2. Further, at 18 and 24 months, three periodicities are significant (*P* < 10^−5^): 1-day, 1/2-day and 1/3-day, indicating the final consolidation of daily sleep-activity rhythms with only one daytime nap in the afternoon. In addition, the proportions of variance explained by the 1/2-day and 1/3-day periodicities in 18 months and 24 months were about the same, both higher compared to 12 months.

**Table 1.** Significant periodicities at 6, 12, 18, 24 months with the corresponding proportions of variances among all Fast Fourier Transform (FFT) signals and p-values.

| Age (Month) | Prop. Variance | Period (min) | Period (day) | p-value |
| --- | --- | --- | --- | --- |
| 6 | 0.0110 | 1440 | 1-day | $6.59 \times 10^{-8}$ |
| 12 | 0.0120 | 1440 | 1-day | $1.75 \times 10^{-8}$ |
| 12 | 0.0068 | 720 | 1/2-day | $3.75 \times 10^{-7}$ |
| 12 | 0.0056 | 360 | 1/4-day | $2.82 \times 10^{-7}$ |
| 12 | 0.0045 | 480 | 1/3-day | $6.49 \times 10^{-6}$ |
| 18 | 0.0130 | 1440 | 1-day | $1.46 \times 10^{-9}$ |
| 18 | 0.0085 | 480 | 1/3-day | $1.87 \times 10^{-10}$ |
| 18 | 0.0083 | 720 | 1/2-day | $5.02 \times 10^{-15}$ |
| 24 | 0.0130 | 1440 | 1-day | $1.97 \times 10^{-9}$ |
| 24 | 0.0086 | 480 | 1/3-day | $1.17 \times 10^{-10}$ |
| 24 | 0.0080 | 720 | 1/2-day | $3.78 \times 10^{-14}$ |

### Comparison with Autocorrelation

To compare the PML algorithm with autocorrelation, the plot of correlation estimates against time lags is shown in Figure 4. The circadian rhythm at 24 hours can be observed at all time points, as the peaks of estimated correlation are at time lags between 23.8 hours to 24.3 hours. We can also observe some local maximal correlation estimates at other time lags: 3.3 hours at 6 months, 4.7 and 10.7 hours at 12 months, 7.5 hours at 18 months, and 7.5 and 16.3 hours at 24 months. While 3.3 hours at 6 months may seem reasonable because of the infant feeding schedule, other cycles are hard to explain [29, 30]. Autocorrelation estimates can be biased due to the presence of multiple periodicities, and generally researchers use it to verify the most dominant periodicity such as 24 hours. Thus from the autocorrelation plots, the most dominant 24-hour rhythm that yields the global maximal correlation estimate can be identified at each age, and this dominant periodicity was also identified by PML.

**Figure 4.**
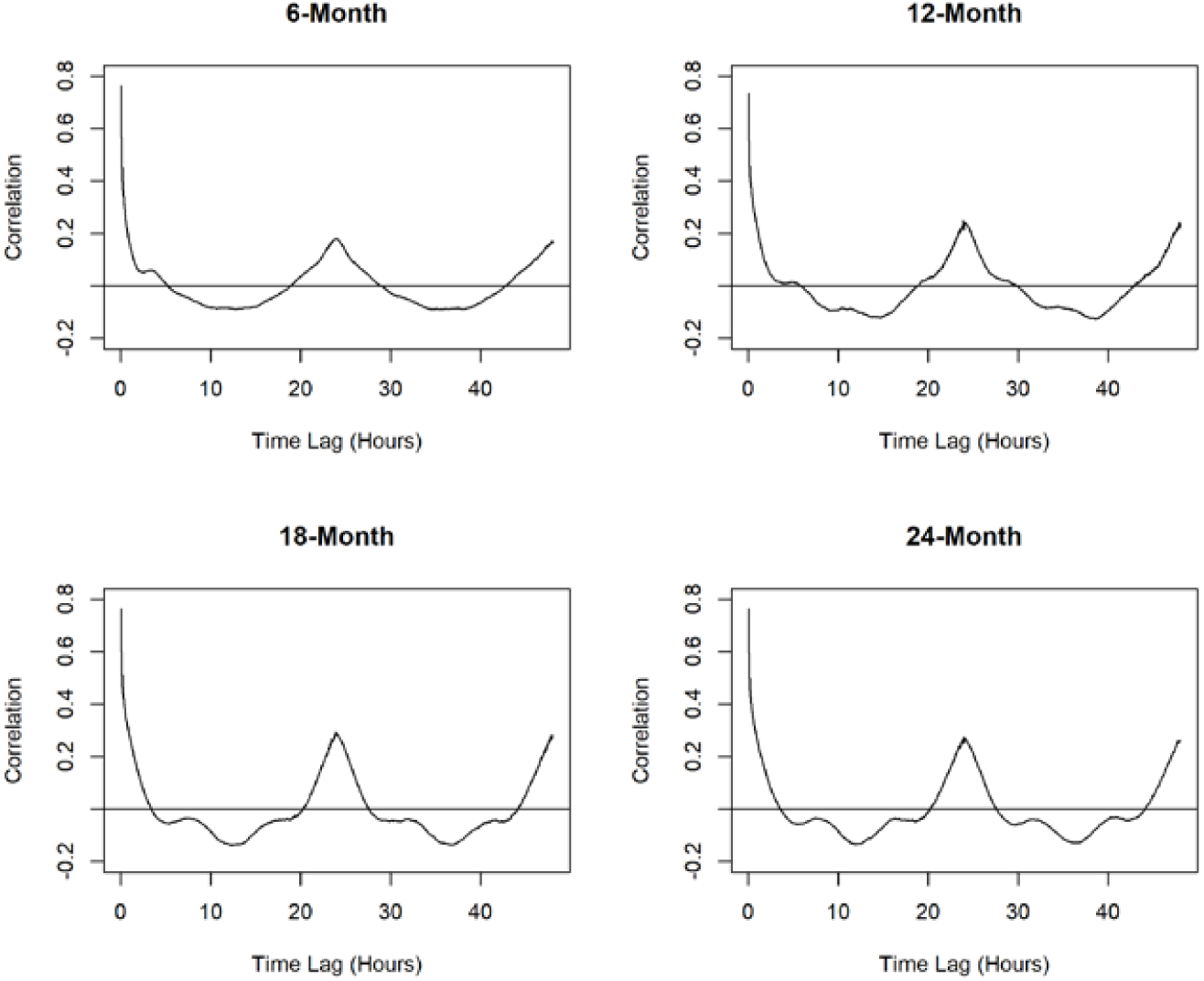
Estimated autocorrelation against time lags at 6, 12, 18, and 24 months respectively.

### Association between Daily Rhythms and Motor Development

Summary of the PDMS-2 standard scores for each category are provided in Table 2.

**Table 2.** The Peabody Developmental Motor Scales-Second Edition (PDMS-2) standard scores for subtests, gross motor, fine motor, and total motor.

| PDMS-2<br>Category | Standard Scores: Mean (Standard Deviation) |  |  |  |
| --- | --- | --- | --- | --- |
|  | 6 Months<br>(n=246) | 12 Months<br>(n=225) | 18 Months<br>(n=192) | 24 Months<br>(n=170) |
| Reflexes | 10.56 (1.02) | -- | -- | -- |
| Stationary | 9.67 (1.44) | 9.48 (1.00) | 9.94 (0.36) | 9.03 (1.38) |
| Locomotor | 10.09 (1.08) | 8.82 (1.76) | 9.26 (1.24) | 8.61 (1.70) |
| Object Manipulation | -- | 9.92 (0.78) | 9.49 (1.16) | 8.74 (1.33) |
| Grasping | 10.69 (1.02) | 11.08 (1.44) | 9.55 (0.78) | 9.99 (0.98) |
| Visual Motor Integration | 11.19 (1.20) | 10.69 (1.13) | 11.04 (1.74) | 9.79 (1.74) |
| Gross Motor | 30.34 (2.72) | 28.25 (2.65) | 28.63 (2.21) | 26.16 (3.17) |
| Fine Motor | 21.86 (2.02) | 21.83 (2.17) | 20.67 (1.93) | 19.80 (2.22) |
| Total | 52.24 (4.18) | 50.23 (5.05) | 49.43 (3.19) | 45.98 (4.69) |

As shown in Table 3, at 6 months, the PDMS-2 total motor scores are found to be associated with assessment seasons (*P* < .001). Infants receiving the PDMS-2 assessment in winter and spring tend to have lower PDMS-2 total motor scores compared to those assessed in summer and autumn. At 12 months, the PDMS-2 total motor scores are associated with both seasons and FFT signals: infants assessed in summer tend to have higher PDMS-2 total motor scores, and infants with higher FFT signals detected at 1/3-day and 1/2-day periodicities also tend to have higher PDMS-2 total motor scores (*P* < .001 and *P*=.04 respectively). *r*^2^ is 0.25 and the adjusted *r*^2^ is 0.21. At 18 and 24 months, no association is identified between the PDMS-2 total motor scores and any other variables.

**Table 3.** Linear regression of Peabody Developmental Motor Scales-Second Edition total motor standard scores on season and Fast Fourier Transform (FFT) signals at 6 months and 12 months, respectively.

| Time | Variable | Estimate | Std. Error | t-value | p-value |
| --- | --- | --- | --- | --- | --- |
| 6-month | (intercept) | 50.10 | 0.56 | 89.52 | <.001 |
|  | spring | 0.48 | 0.85 | 0.56 | .576 |
|  | summer | 4.48 | 0.82 | 5.43 | <.001 |
|  | autumn | 4.06 | 0.89 | 4.54 | <.001 |
| 12-month | (intercept) | 45.52 | 1.29 | 35.25 | <.001 |
|  | summer | 1.66 | 1.58 | 1.05 | .297 |
|  | 1/2-day* | 0.19 | 0.09 | 2.10 | .039 |
|  | 1/3-day* | 0.31 | 0.09 | 3.55 | <.001 |
|  | summer:1/3-day* | -0.48 | 0.16 | -3.00 | .004 |

\* FFT signals were multiplied by 10,000 in regression models so that the estimated effect sizes are for every 10,000-unit increase in FFT signals.

Because PDMS-2 total motor scores are associated with FFT signals at 12 months, further linear regression between each subtest scores and FFT signals are also examined. As shown in Table 4, subtests for stationary and locomotion as well as gross motor and fine motor are found to be associated with the 1/3-day periodicity. Gross motor represents the overall performance on the three subtests of stationary, locomotion and object manipulation for infants at 12 months, and since the association of FFT signals at 1/3-day periodicity with stationary and locomotion subtests is strong, it is expected that the association of that with gross motor is also strong. *r*^2^ and the adjusted *r*^2^ are 0.05 and 0.04 for the stationary model, 0.23 and 0.20 for the locomotion model, 0.21 and 0.17 for the gross motor model, and 0.15 and 0.11 for the fine motor model respectively.

**Table 4.** Linear regression of Peabody Developmental Motor Scales-Second Edition standard scores on season and Fast Fourier Transform (FFT) signals at 12 months: stationary and locomotion subtests, gross motor and fine motor as the outcome respectively.

| Category | Variable | Estimate | Std. Error | t-value | p-value |
| --- | --- | --- | --- | --- | --- |
| Stationary | (Intercept) | 9.23 | 0.19 | 47.9 | <.001 |
|  | 1/3-day | 0.04 | 0.02 | 2.17 | .033<sup>a</sup> |
| Locomotion | (Intercept) | 7.16 | 0.43 | 16.48 | <.001 |
|  | summer | 0.71 | 0.75 | 0.94 | .348 |
|  | 1/3-day | 0.19 | 0.04 | 4.61 | <.001<sup>a</sup> |
|  | summer: 1/3-day | -0.13 | 0.07 | -1.93 | .057<sup>b</sup> |
| Gross Motor | (Intercept) | 26.41 | 0.63 | 42.03 | <.001 |
|  | summer | 0.57 | 1.09 | 0.52 | .602 |
|  | 1/3-day | 0.24 | 0.06 | 3.97 | <.001<sup>a</sup> |
|  | summer: 1/3-day | -0.17 | 0.10 | -1.73 | .087<sup>b</sup> |
| Fine Motor | (Intercept) | 21.08 | 0.5 | 42.15 | <.001 |
|  | summer | 0.34 | 0.86 | 0.39 | .697 |
|  | 1/3-day | 0.11 | 0.05 | 2.36 | .021<sup>a</sup> |
|  | summer: 1/3-day | -0.15 | 0.08 | -1.89 | .062<sup>b</sup> |

\* FFT signals were multiplied by 10,000 in regression models so that the estimated effect sizes are for every 10,000-unit increase in FFT signals.
<sup>a</sup> Statistical significance level at $\alpha=0.05$ .
<sup>b</sup> Statistical significance level at $\alpha=0.10$ .

## Discussion

### Method Evaluation

The PML approach is very effective in studying daily activity rhythms among infants and toddlers. At 6 and 12 months, the dominant 1-day periodicity suggests the formation of the 24-hour cycle. At 18 and 24 months, the combination of the dominant 1-day, 1/3-day, and 1/2-day periodicities forms the consolidated daily activity pattern with two activity peaks during the day and one afternoon nap. PML not only effectively identified population-level dominant periodicities but also characterized sleep-activity patterns without complex functional analysis. PML can complement current methods for circadian rhythm analysis and is especially useful for populations whose daily rhythm patterns are non-sinusoidal and irregular. On the other hand, because PML is applicable to time series data bearing a similar nature to actigraphy, the application of PML can be extended to other types of circadian rhythm studies using information such as body temperature and hormone data to study and characterize daily rhythms effectively.

### Comparison with Other Methods

In comparison, Fisher’s test for harmonic analysis tends to identify many significant periodicities unless a stringent threshold is used for statistical significance. Here we employed the Bonferroni correction to adjust for multiple testing and used the significance level 10^−5^ to select periodicities, even though we did not conduct as many statistical tests simultaneously. In sequential testing procedures like our scenario, people often use less conservative multiple testing correction methods such as the Benjamini-Hochberg procedure [31]. We chose the most stringent threshold to avoid selecting too many periodicities that are not helpful in characterizing daily activity patterns at each age.

We also compare our PML algorithm with the standard approach autocorrelation. Plots of correlation estimates against time lags are useful for identifying the correlation peak at 24 hours but not shorter periods of rhythmicity. It is because estimation of correlation can be biased due to the presence of multiple periodic rhythms, and identification of multiple periodicities by simple calculation of autocorrelation may not be accurate. Therefore, the standard approach using autocorrelation is effective in confirming the most dominant 24-hour periodicity but not as effective in identifying other periodicities, which the PML algorithm is able to achieve.

Other standard approaches such as periodograms and cosinor analysis are not used here, because there are in fact connections between PML and the two methods. It is important to point out their connections and also differences. For periodograms, the Fourier periodogram, the Enright periodogram and the Lomb-Scargle periodogram are commonly used [18]. Both our PML algorithm and the Fourier periodogram utilize Fourier analysis to identify dominant periodicities, except that the PML algorithm uses a shrinkage method in machine learning to identify dominant patterns in the population, while Fourier periodograms are focused on individuals to manually identify dominant periodicities based on individual plots. The Enright periodogram, though suitable for equidistant activity measurements in our scenario, may not be applicable here because it requires ten or more days of data [18]. In addition, the estimation method only holds when there is one periodic component, but in our case the presence of multiple periodic components may attenuate the results [18]. The Lomb-Scargle periodogram is a modification of Fourier analysis to accommodate unevenly spaced data or missing data. Because our data do not have the issue, the Lomb-Scargle periodogram is equivalent to the Fourier periodogram here. Compared with the PML algorithm, the Fourier periodogram involves more manual work to generate periodograms for each individual and visually identify dominant periodicities, while PML is more automated and also more effective in studying the population as a whole and further identifying the periodicities that are characteristic of the population. In addition, researchers often use periodograms to validate the most dominant periodicity such as 24 hours but do not specifically examine information on secondary dominant periodicities or use periodic curves to reconstruct or approximate activity patterns, even though the connection between dominant frequencies/periodicities and functional curves can be made and the periodic information can be fully utilized. Therefore, the PML algorithm makes full use of the information from more than one dominant periodicities and links FFT results in the frequency domain with their representing cosine curves to effectively characterize activity patterns.

For cosinor analysis, recall that FFT results consist of real parts and imaginary parts that correspond to cosine curves and sine curves respectively, and thus FFT is equivalent to fitting the cosine model. We actually fitted cosine models to the activity data with one to three cosine curves at dominant periodicities identified by the PML algorithm. Even though the estimated amplitudes for the cosine curves are different from FFT results, the Pearson correlation between the cosine coefficients and the FFT signals of the same periodicity is 1, indicating equivalence. While the final results are equivalent, the procedures are different. For cosinor analysis, if prior knowledge is known, one can use simple least squares methods to fit the model [32]. However, if there is no prior knowledge on periodic information, the least squares methods cannot be used because the dominant periodicity needs to be estimated first and the cosinor model can no longer be linearized in its parameters. One has to either start from an initial guess and use iterative procedures to minimize residual sum of squares or use simulated annealing alternatively to fit the model, the process of which can be exhaustive [33-35]. In comparison, without prior knowledge on dominant periods, the PML algorithm based on shrinkage in machine learning is still easy to implement without computational burden in extracting periodic information and the results are as effective as the cosinor model to characterize daily activity patterns using cosine curves.

In summary, the proposed PML algorithm is effective in extracting periodic information, identifying dominant periodicities, and further characterizing activity patterns. In the presence of multiple periodicities, PML does not have the estimation problem that autocorrelation encounters. To identify dominant periodicities, PML uses shrinkage in machine learning methods that can avoid manual work in periodograms, which require individual plots and visual identification. PML can also characterize activity patterns by making full use of the cosine curves represented by FFT signals and avoiding the computationally intensive process of choosing and fitting cosinor models when prior knowledge on dominant periodicities is not available.

### Sleep-Activity Rhythm Characterization

Our study confirms previous study results that infants already form 24-hour sleep-wake cycles at 6 months due to entrainment by cyclical changes in the environment, whether it is due to light-dark cycles or maternal rest-activity cycles [6, 9, 10, 36, 37]. It is noteworthy that while 24-hour cycles are formed, sleep-activity patterns over the 24-hour period are not stabilized: infants often take multiple naps at different times of the day and their daily activity patterns vary across days and across individuals.

From 6 months to 12 months, our study indicates that the infant sleep-activity pattern gradually develops: some infants take only one afternoon nap while others take two naps in the morning and afternoon. Strong FFT signals at 1/3-day periodicity capture two-peak one-nap activity patterns while strong FFT signals at 1/4-day periodicity capture three-peak two-nap patterns. The results are in line with previous sleep studies that the duration of nighttime sleep gradually lengthens and sleep patterns become more and more consolidated during the first year after birth [12, 38].

While the timing for the stabilization of infant sleep-activity patterns varies across individuals, by the time infants become 18 months old, their daily activity patterns have consolidated into a predominant nighttime pattern with one afternoon nap only, which can be obtained by combining the three dominant periodicities at 1-day, 1/2-day and 1/3-day. The consolidation of sleep-activity patterns is confirmed by increased FFT signals at 1/2-day and 1/3-day periodicities and decreased FFT signals at other periodicities compared to previous ages. The results for 24 months remain the same as 18 months, showing no changes and confirming further that sleep-activity patterns are consolidated by 18 months and stable onwards.

In our study, the 3-hour periodicity, normally for feeding behaviors, was not detected and there might be two reasons. First, in the feeding guidelines for infants, feeding of infants aged 6-8 months are advised to be 5 to 6 times in 24 hours, less frequent than the 3-hour (8 times) schedule, and infants aged 12-24 months are advised to have three meals with family and have additional snacks for 2-3 times [29, 30]. Infants younger than 6 months may have a more frequent feeding schedule, but our activity data were collected at 6 months or later. Second, there might be desynchronization between feeding schedules and activity patterns. While 6-month infants might be fed 5-6 times per day, they do not nap or sleep 5-6 times within the same timeframe. We referred to the sleep diaries recorded by parents as reference for napping information. Most 6-month infants have one to two naps in the morning and one nap in the afternoon. 12-month infants generally have one or no naps in the morning and one nap in the afternoon. Most 18-month and 24-month infants have one nap in the afternoon. Therefore, sleep-activity patterns are desynchronized with feeding schedules, as feeding behaviours might not dominate infant sleep-activity patterns at this age. For the above reasons, feeding cycles such as 3-hour periodicity were not detected in our data.

### Association between Daily Rhythms and Early Childhood Development

Using FFT signals at dominant periodicities identified by PML, we are able to find the association between the consolidation of sleep-activity rhythms and early childhood motor development. At 6 months, the association between PDMS-2 total motor scores and assessment seasons may be explained by different amounts of clothes worn by infants in different seasons. In winter, infants are likely to wear lots of clothes, which may restrict their behaviors and result in suboptimal performances compared to infants wearing light clothes and taking the PDMS-2 assessment in summer. As a result, 6-month-old infants taking the assessment in summer and autumn got relatively higher PDMS-2 scores compared to infants in winter and spring.

Then at 12 months, after controlling for assessment seasons, stronger FFT signals at 1/2-day and 1/3-day periodicities are generally associated with higher PDMS-2 scores. 12 months is in the critical period of sleep-activity rhythm consolidation, which is captured by the growing FFT signals at 1/2-day and 1/3-day periodicities. It is noteworthy that for infants at this age, they all have strong FFT signals at 1-day periodicity, indicating that they all exhibit 1-day periodicity in their sleep-activity patterns and their 24-hour periodic activity patterns tend to be stabilized. As a result, there is not much variation in the strength of 1-day periodicity, which might not explain much of the variability in PDMS-2 scores among inidividuals. In comparison, the activity pattern over the 24-hour period is not consolidated and the activity pattern can be characterized by the 1/3-day and 1/2-day periodicities. The larger variability in the strength of 1/3-day and 1/2-day periodicities can describe the degree to which the daily sleep-activity pattern is consolidated, which is further associated with children development evaluated by PDMS-2 scores. Infants with a more consolidated activity pattern tend to have better motor development evaluations. In addition, activity rhythm consolidation is strongly associated with subcategories of locomotion and stationary, which belong to the gross motor and measure how the large muscle system is utilized to move from place to place or assume a stable posture when not moving. Therefore, we obtained new insights into early childhood development that the degree to which the sleep-activity pattern is consolidated at 12 months is associated with infant motor development and is associated with the large muscle system development in particular.

At 18 and 24 months, the PDMS-2 scores are not associated with season, FFT signals, or any other variables in the dataset we have. Most of the toddlers have stabilized daily activity patterns with strong periodic rhythmicity at this age. FFT signals as characteristics of sleep-activity rhythms are no longer associated with PDMS-2 scores, and it is likely because the critical age at which the daily activity rhythm stabilizes has passed.

### Limitations and Future Work

One limitation with our study is that we collected Actiwatch data every six months, and thus we were not able to capture more detailed monthly changes over the time period. Future work may collect Actiwatch data in a more frequent manner, such as every three months or every month, so as to capture gradual changes in the sleep-activity rhythm during early childhood. For infants younger than 6 months, more frequent observations can also allow us to observe how the predominant rhythm of infants changes from ultradian to circadian by adjusting to the 24-hour cycle in the environment. Another limitation of the study is that while we identified the association of sleep-activity daily rhythm consolidation with early childhood motor development and with the large muscle development in particular, the mechanism behind it is not clear. Future work should investigate how daily rhythm consolidation and motor development interrelate and contribute to early childhood development.

## Conclusions

In summary, the proposed PML algorithm provides a new method for circadian rhythm analysis and is particularly useful for studying populations whose daily patterns are not regular. In addition, the PML algorithm is applicable to other types of wearable device data in the format of time series bearing a similar nature to actigraphy, so it can be extended to other types of circadian studies using information such as body temperature, heart rate, and hormone data. Therefore, the PML algorithm can be widely applied to other wearable device studies to help characterize periodic information. Using the proposed method, our study provides novel insights into sleep-activity rhythm development in early childhood. First, in our study, the critical time period for the consolidation of sleep-activity rhythms is between 6 and 18 months. It is because at 6 months, 24-hour sleep-wake cycles are formed but their daily activity patterns are not stabilized, and by the time toddlers become 18 months old, their sleep-activity patterns have consolidated into a fixed pattern with two activity peaks and one afternoon nap. The time period between 6 and 18 months is critical for early childhood sleep-activity rhythm development and consolidation. Second, we identified the association between the consolidation of daily rhythms and early childhood motor development, and the association is with the large muscle system development in particular. This association has not been identified in previous studies. Infants with more consolidated circadian rhythms tend to have better motor development assessments. While the mechanism is not clear, keeping a regular and stable sleep-activity pattern and maintaining a healthy circadian system are important for healthy physical development in early childhood.

## Funding

The study was supported by the Chinese National Natural Science Foundation (81773443); Ministry of Science and Technology of China (2016YFC1305203); Ministry of Health of the People’s Republic of China (201002006); Shanghai Science and Technology Commission (17XD1402800, 18695840200); Construction Project of Key and Weak Discipline of Shanghai municipal Commission of Health and Family Planning (2016ZB0104); Shanghai Jiao Tong University (YG2016ZD04).

## Acknowledgments

We would like to thank all parents and children who participated in the study.

## Conflict of Interests

The authors have no potential conflicts of interest relevant to this article to disclose.

## Abbreviations

FFT: Fast Fourier Transform
MSE: mean squared error
PDMS-2: Peabody Developmental Motor Scales-Second Edition
PML: Penalized Multi-band Learning
PSG: polysomnography

## Notes

### Competing Interest Statement

The authors have declared no competing interest.

### Summary of Updates

The title of the manuscript

